# ExoNet Database: Wearable Camera Images of Human Locomotion Environments

**DOI:** 10.1101/2020.10.23.352054

**Authors:** Brock Laschowski, William McNally, Alexander Wong, John McPhee

## Abstract

Advances in computer vision and artificial intelligence are allowing researchers to develop environment recognition systems for powered lower-limb exoskeletons and prostheses. However, small-scale and private training datasets have impeded the widespread development and dissemination of image classification algorithms for classifying human walking environments. To address these limitations, we developed ExoNet - the first open-source, large-scale hierarchical database of high-resolution wearable camera images of human locomotion environments. Unparalleled in scale and diversity, ExoNet contains over 5.6 million RGB images of different indoor and outdoor real-world walking environments, which were collected using a lightweight wearable camera system throughout the summer, fall, and winter seasons. Approximately 923,000 images in ExoNet were human-annotated using a 12-class hierarchical labelling architecture. Available publicly through IEEE DataPort, ExoNet offers an unprecedented communal platform to train, develop, and compare next-generation image classification algorithms for human locomotion environment recognition. Besides the control of powered lower-limb exoskeletons and prostheses, applications of ExoNet could extend to humanoids and autonomous legged robots.

## 1 Introduction

Hundreds of millions of individuals worldwide have mobility impairments resulting from degenerative aging and neuro-musculoskeletal disorders like spinal cord injury, osteoarthritis, Parkinson’s disease, and cerebral palsy (Grimmer et al., 2019). Fortunately, newly-developed powered lower-limb exoskeletons and prostheses can allow otherwise wheelchair-bound seniors and rehabilitation patients to perform movements that involve net positive mechanical work (e.g., climbing stairs and standing from a seated position) using onboard actuators and intelligent control systems (Krausz and Hargrove, 2019; Laschowski and Andrysek, 2018; Tucker et al., 2015; Young and Ferris, 2017; Zhang et al., 2019a). Generally speaking, the high-level controller recognizes the patient’s locomotion mode (intention) by analyzing real-time measurements from wearable sensors using machine learning algorithms. The mid-level controller then translates the locomotion intentions into mode-specific reference trajectories. This control level typically comprises a finite state machine, which implements a discrete parametrized control law (e.g., joint position or mechanical impedance control) for each different locomotion mode. Finally, the low-level controller tracks the reference trajectories and minimizes the signal error by modulating the device actuators using feedforward and feedback control loops (Krausz and Hargrove, 2019; Laschowski and Andrysek, 2018; Tucker et al., 2015; Young and Ferris, 2017; Zhang et al., 2019a).

Accurate transitions between different locomotion modes is important since even rare misclassifications can cause loss-of-balance and injury. In many commercial devices like the ReWalk and Indego powered lower-limb exoskeletons, the patient acts as the high-level controller by performing volitional movements to manually switch between locomotion modes (Tucker et al., 2015; Young and Ferris, 2017). These human-controlled methods can be time-consuming, inconvenient, and cognitively demanding. Researchers have recently developed automated locomotion mode recognition systems using wearable sensors like inertial measurement units (IMUs) and surface electromyography (EMG) to automatically switch between different locomotion modes (Krausz and Hargrove, 2019; Laschowski and Andrysek, 2018; Tucker et al., 2015; Young and Ferris, 2017; Zhang et al., 2019a). Whereas mechanical and inertial sensors respond to the patient’s movements, the electrical potentials of biological muscles, as recorded using surface EMG, precede movement initiation and thus could (marginally) predict locomotion mode transitions. Several researchers have combined mechanical sensors with surface EMG for automated locomotion mode recognition. Such neuromuscular-mechanical data fusion has improved the locomotion mode recognition accuracies and decision times compared to implementing either system individually (Du et al., 2012; Huang et al., 2011; Liu et al., 2016; Wang et al., 2013). However, these measurements are still patient-dependent, and surface EMG are susceptible to fatigue, changes in electrode-skin conductivity, and crosstalk from adjacent muscles (Tucker et al., 2015).

Supplementing neuromuscular-mechanical data with information about the upcoming walking environment could improve the high-level control performance. Analogous to the human visual system, environment sensing would precede modulation of the patient’s muscle activations and/or walking biomechanics, therein enabling more accurate and real-time locomotion mode transitions. Environment sensing can also be used to adapt low-level reference trajectories (e.g., changing toe clearance corresponding to an obstacle height) (Zhang et al., 2020) and optimal path planning (e.g., identifying opportunities for energy regeneration) (Laschowski et al., 2019a; 2020a). Preliminary research has shown that supplementing a locomotion mode recognition system with environment information can improve the classification accuracies and decision times compared to excluding terrain information (Huang et al., 2011; Liu et al., 2016; Wang et al., 2013). Several researchers have explored using radar detectors (Kleiner et al., 2018) and laser rangefinders (Liu et al., 2016; Wang et al., 2013; Zhang et al., 2011) for environment sensing. However, wearable vision-based systems can provide more detailed information about the field-of-view and detect physical obstacles in peripheral locations. Most environment recognition systems have included either RGB cameras (Da Silva et al., 2020; Diaz et al., 2018; Khademi and Simon, 2019; Krausz and Hargrove, 2015; Laschowski et al., 2019b; Novo-Torres et al., 2019; Zhong et al., 2020) or 3D depth cameras (Hu et al., 2018; Krausz et al., 2015; 2019; Massalin et al., 2018; Varol and Massalin, 2016; Zhang et al., 2019b; 2019c; 2019d).

For image classification, researchers have used learning-based algorithms like support vector machines (Massalin et al., 2018; Varol and Massalin, 2016) and deep convolutional neural networks (Khademi and Simon, 2019; Laschowski et al., 2019b; Novo-Torres et al., 2019; Rai and Rombokas, 2018; Zhang et al., 2019b; 2019c; 2019d; Zhong et al., 2020). Although convolutional neural networks typically outperform support vector machines for image classification (LeCun et al., 2015), deep learning requires significant and diverse training images to prevent overfitting and promote generalization. Deep learning has become pervasive ever since AlexNet (Krizhevsky et al., 2012) popularized convolutional neural networks by winning the 2012 ImageNet challenge. ImageNet is an open-source dataset containing ~15 million labelled images and 22,000 different classes (Deng et al., 2009). The lack of an open-source, large-scale research dataset of human locomotion environment images has impeded the development of environment-aware control systems for powered lower-limb exoskeletons and prostheses. Until now, researchers have been required to individually collect training images to develop their classification algorithms. These repetitive measurements are time-consuming and inefficient, and individual private datasets have prevented comparisons between classification algorithms from different researchers (Laschowski et al., 2020b). Drawing inspiration from ImageNet, we developed ExoNet - the first open-source, large-scale hierarchical database of high-resolution wearable camera images of human walking environments. In accordance with the Frontiers submission guidelines, this article provides a detailed description of the research dataset. Benchmark performance and analyses of the ExoNet database for human locomotion environment classification will be presented in future work.

## 2 Materials and Methods

### 2.1 Large-Scale Data Collection

One subject was instrumented with a lightweight wearable smartphone camera system (iPhone XS Max); photograph shown in Figure 1A. Unlike limb-mounted systems (Da Silva et al., 2020; Diaz et al., 2018; Hu et al., 2018; Kleiner et al., 2018; Massalin et al., 2018; Rai and Rombokas, 2018; Varol and Massalin, 2016; Zhang et al., 2011; 2019b; 2019c), chest-mounting can provide more stable video recording and allow users to wear pants and long dresses without obstructing the sampled field-of-view. The chest-mount height was approximately 1.3 m from the ground when the participant stood upright. The smartphone contains two 12-megapixel RGB rear-facing cameras and one 7-megapixel front-facing camera. The front and rear cameras provide 1920×1080 and 1280×720 video recording at 30 frames/second, respectively. The smartphone weighs approximately 0.21 kg, and features an onboard rechargeable lithium-ion battery, 512-GB of memory storage, and a 64-bit ARM-based integrated circuit (Apple A12 Bionic) with six-core CPU and four-core GPU. These hardware specifications can support onboard machine learning for real-time environment classification. The relatively lightweight and unobtrusive nature of the wearable camera system allowed for unimpeded human walking biomechanics. Ethical review and approval were not required for this research in accordance with the University of Waterloo Office of Research Ethics.

**Figure 1.**
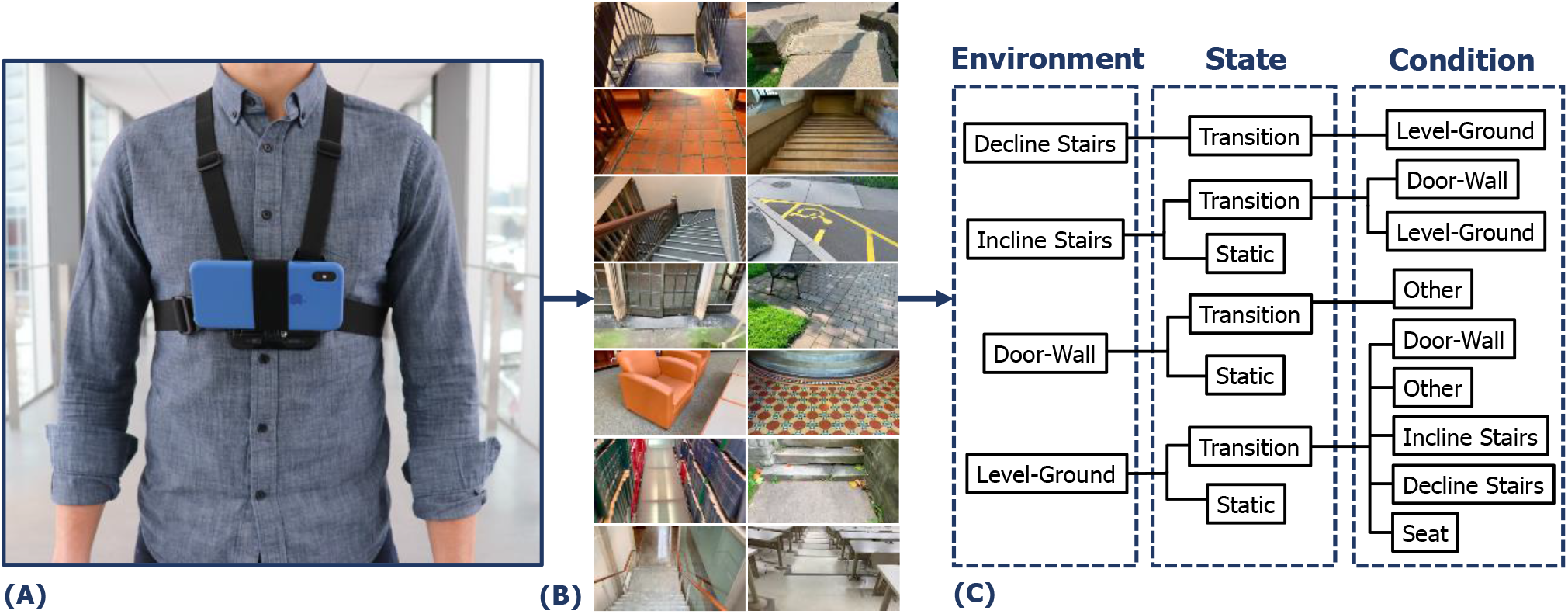
Development of the ExoNet database, including **(A)** photograph of the wearable camera system used for large-scale data collection; **(B)** examples of the high-resolution RGB images (1280×720) of human walking environments; and **(C)** schematic of the 12-class hierarchical labelling architecture.

While most environment recognition systems have been limited to controlled indoor environments and/or prearranged walking circuits (Du et al., 2012; Hu et al., 2018; Khademi and Simon, 2019; Kleiner et al., 2018; Krausz et al., 2015; 2019; Liu et al., 2016; Wang et al., 2013; Zhang et al., 2011; 2019b; 2019c; 2019d), our subject walked around unknown outdoor and indoor real-world environments while collecting images with occlusions, signal noise, and intraclass variations. Data were collected at various times throughout the day to incorporate different lighting conditions. Analogous to human gaze fixation during walking (Li et al., 2019), the sampled field-of-view was approximately 1-5 meters ahead of the participant, thereby showing upcoming walking environments rather than the ground underneath the subject’s feet. The camera’s pitch angle slightly differed between data collection sessions. Images were sampled at 30 Hz with 1280×720 resolution. More than 52 hours of video were recorded, amounting to approximately 5.6 million total images (examples shown in Figure 1B). The same environment was never sampled twice to maximize diversity among the ExoNet images. Data were collected throughout the summer, fall, and winter seasons to incorporate different weathered surfaces like snow, grass, and multicolored leaves. In accordance with the Frontiers submission guidelines, the ExoNet database was deposited in a public repository (IEEE DataPort) and is available for download at https://ieee-dataport.org/open-access/exonet-database-wearable-camera-images-human-locomotion-environments. The file size of the uncompressed videos is approximately 140 GB.

### 2.2 Hierarchical Image Labelling

Given the subject’s preferred walking speed, there were minimal differences between consecutive images sampled at 30 Hz. The labelled images were therefore downsampled to 5 frames/second to minimize the demands of manual annotation and increase the diversity in image appearances. However, for real-time environment classification and control of powered lower-limb exoskeletons and prostheses, higher sampling rates would be more advantageous for accurate locomotion mode recognition and transitioning. Similar to ImageNet (Deng et al., 2009), the ExoNet database was human-annotated using a hierarchical labelling architecture (see Figure 1C). Images were labelled according to exoskeleton and prosthesis control functionality, rather than a purely computer vision perspective. For instance, images of level-ground environments showing either pavement or grass were not differentiated since both surfaces would use the same level-ground walking state controller. In contrast, computer vision researchers might label these different surface textures as separate classes.

Approximately 923,000 images in ExoNet were manually labelled and organized into 12 classes using the following descriptions, which also include the number of labelled images/class: {IS-T-DW = 31,628} shows incline stairs with a door and/or wall; {IS-T-LG = 11,040} shows incline stairs with level-ground thereafter; {IS-S = 17,358} shows only incline stairs; {DS-T-LG = 28,677} shows decline stairs with level-ground thereafter; {DW-T-O = 19,150} shows a door and/or wall with *other* (e.g., hand or window); {DW-S = 36,710} shows only a door and/or wall; {LG-T-DW = 379,199} shows level-ground with a door and/or wall; {LG-T-O = 153,263} shows level-ground with *other* (e.g., humans, cars, bicycles, or garbage cans); {LG-T-IS = 26,067} shows level-ground with incline stairs thereafter; {LG-T-DS = 22,607} shows level-ground with decline stairs thereafter; {LG-T-SE = 119,515} shows level-ground with seats (e.g., couches, chairs, or benches); and {LG-S = 77,576} shows only level-ground. These classes were selected post hoc to encompass the different walking environments encountered during the data collection sessions. We included the *other* class to improve image classification performance when confronted with non-terrain related features like humans and bicycles.

Inspired by previous work (Du et al., 2012; Huang et al., 2011; Khademi and Simon, 2019; Liu et al., 2016; Wang et al., 2013), the hierarchical labelling architecture included both *static* (S) and *transition* (T) states. A static state describes an environment where an exoskeleton or prosthesis user would continuously perform the same locomotion mode (e.g., only level-ground terrain). In contrast, a transition state describes an environment where the exoskeleton or prosthesis high-level controller might switch between different locomotion modes (e.g., level-ground and incline stairs). Manually labelling the transition states was relatively subjective. For example, an image showing level-ground terrain was labelled *level-ground-transition-incline stairs* (LG-T-IS) when an incline staircase was approximately within the sampled field-of-view and forward-facing. Similar labelling principles were applied to transitions to other conditions. The Python code used for labelling the ExoNet database was uploaded to GitHub and is publicly available for download at https://github.com/BrockLaschowski2/ExoNet.

## 3 Discussion

Environment recognition can improve the control of powered lower-limb exoskeletons and prostheses during human locomotion. However, small-scale and private training datasets have impeded the widespread development and dissemination of image classification algorithms for human locomotion environment recognition. Motivated by these limitations, we developed ExoNet - the first open-source, large-scale hierarchical database of high-resolution wearable camera images of human walking environments. Using a lightweight wearable camera system, we collected over 5.6 million RGB images of different indoor and outdoor real-world walking environments, of which approximately 923,000 images were human-annotated using a 12-class hierarchical labelling architecture. Available publicly through IEEE DataPort, ExoNet provides researchers an unprecedented communal platform to develop and compare next-generation image classification algorithms for human locomotion environment recognition. Although ExoNet was originally designed for environment-aware control of powered lower-limb exoskeletons and prostheses, applications could extend to humanoids and autonomous legged robots (Park et al., 2015; Villarreal et al., 2020). Users of the ExoNet database are requested to reference this dataset report.

Aside from being the only open-source database of human locomotion environment images, the scale and diversity of ExoNet significantly distinguishes itself from previous environment recognition systems, as illustrated in Table 1. ExoNet contains approximately 923,000 labelled images. In comparison, the previous largest dataset contained approximately 402,000 images (Massalin et al., 2018). While most environment recognition systems have included fewer than 6 classes (Khademi and Simon, 2019; Krausz and Hargrove, 2015; Krausz et al., 2015; 2019; Laschowski et al., 2019b; Massalin et al., 2018; Novo-Torres et al., 2019; Varol and Massalin, 2016; Zhang et al., 2019b; 2019c; 2019d; 2020), the ExoNet database features a 12-class hierarchical labelling architecture. These differences have practical implications given that learning-based algorithms like deep convolutional neural networks require significant and diverse training images (LeCun et al., 2015). The spatial resolution of the ExoNet images (1280×720) is considerably higher than previous efforts (e.g., 224×224 and 320×240). Poor image resolution has been attributed to decreased classification accuracy of human locomotion environments (Novo-Torres et al., 2019). Although higher resolution images can increase the computational and memory storage requirements, that being unfavourable for real-time mobile computing, research has been moving towards the development of efficient convolutional neural networks that require fewer operations (Tan and Le, 2020), therein enabling the processing of larger images for relatively similar computational power. Here we assume mobile computing for exoskeleton and prosthesis control (i.e., untethered and no wireless communication to cloud computing). Nevertheless, an exoskeleton or prosthesis controller may not always benefit from additional information provided by higher resolution images, particularly when interacting with single surface textures (i.e., only pavement or grass). With ongoing research and development in computer vision and artificial intelligence, larger and more challenging training datasets are needed to develop better image classification algorithms for environment-aware locomotor control systems.

**Table 1.** Comparison of the ExoNet database with previous environment recognition systems for powered lower-limb prostheses and exoskeletons.

| Reference | Sensor | Position | Dataset | Resolution | Classes |
| --- | --- | --- | --- | --- | --- |
| Da Silva et al (2020) | RGB Camera | Lower-Limb | 3,992 Images | 512×512 | 6 |
| Diaz et al (2018) | RGB Camera | Lower-Limb | 3,992 Images | 1080x1920 | 6 |
| Khademi and Simon (2019) | RGB Camera | Waist | 7,284 Images | 224x224 | 3 |
| Krausz and Hargrove (2015) | RGB Camera | Head | 5 Images | 928x620 | 2 |
| Krausz et al (2015) | Depth Camera | Chest | 170 Images | 80x60 | 2 |
| Krausz et al (2019) | Depth Camera | Waist | 4,000 Images | 171x224 | 5 |
| Laschowski et al (2019b) | RGB Camera | Chest | 34,254 Images | 224x224 | 3 |
| Massalin et al (2018) | Depth Camera | Lower-Limb | 402,403 Images | 320x240 | 5 |
| Novo-Torres et al (2019) | RGB Camera | Head | 40,743 Images | 128x128 | 2 |
| Varol and Massalin (2016) | Depth Camera | Lower-Limb | 22,932 Images | 320x240 | 5 |
| Zhang et al (2019b; 2019c) | Depth Camera | Lower-Limb | 7,500 Images | 224x171 | 5 |
| Zhang et al (2019d) | Depth Camera | Waist | 4,016 Images | 2048 Point Cloud | 3 |
| Zhang et al (2020) | Depth Camera | Lower-Limb | 7,500 Images | 100x100 | 5 |
| Zhong et al (2020) | RGB Camera | Head and Lower-Limb | 327,000 Images | 1240x1080 | 6 |
| **ExoNet Database** | **RGB Camera** | **Chest** | **922,790 Images** | **1280×720** | **12** |

A potential limitation of the ExoNet database is the two-dimensional nature of the environment information. Whereas RGB cameras measure light intensity information, depth cameras also provide distance measurements (Hu et al., 2018; Krausz et al., 2015; 2019; Massalin et al., 2018; Varol and Massalin, 2016; Zhang et al., 2019b; 2019c; 2019d). Depth cameras work by emitting infrared light and calculate distance by measuring the light time-of-flight between the camera and physical environment (Varol and Massalin, 2016). Depth measurement accuracies typically degrade in outdoor lighting conditions (e.g., sunlight) and with increasing measurement distance. Consequently, most environment recognition systems using depth cameras have been tested in indoor environments (Hu et al., 2018; Krausz et al., 2015; 2019; Massalin et al., 2018; Varol and Massalin, 2016) and have had limited capture volumes (i.e., between 1-2 m of maximum range imaging) (Krausz et al., 2015; Massalin et al., 2018; Varol and Massalin, 2016). Moreover, assuming mobile computing, the application of depth cameras for environment sensing would require powered lower-limb exoskeletons and prostheses to have embedded microcomputers with significant computing power and minimal power consumption, the specifications of which are not supported by existing untethered systems (Massalin et al., 2018). These practical limitations motivated our decision to use RGB images.

The wearable camera images could be fused with the smartphone IMU measurements to improve high-level control performance. For example, if an exoskeleton or prosthesis user unexpectedly stops while walking towards an incline staircase, the acceleration measurements would indicate static standing rather than stair ascent, despite the staircase being accurately detected in the field-of-view. Since environment information does not explicitly represent the locomotor intent, environment recognition systems should supplement, rather than replace, automated locomotion mode recognition systems based on patient-dependant measurements like mechanical and inertial sensors. The smartphone IMU measurements could also be used for sampling rate control (Da Silva et al., 2020; Diaz et al., 2018; Khademi and Simon, 2019; Zhang et al., 2011). Faster walking speeds would likely benefit from higher sampling rates for continuous classification. In contrast, static standing does not necessarily require environment information and therefore the smartphone camera could be powered down, or the sampling rate decreased, to minimize the computational and memory storage requirements. However, the optimal method for fusing the smartphone camera images with the onboard IMU measurements remains to be determined.

## 5 Author Contributions

BL was responsible for the study design, literature review, data collection, image labelling, data interpretation, and manuscript writing. WM assisted with the study design, image labelling, data interpretation, and manuscript writing. AW and JM assisted with the study design, data interpretation, and manuscript writing. All authors read and approved the final manuscript.

